# Evolution to alternative levels of stable diversity leaves areas of niche space unexplored

**DOI:** 10.1101/2021.01.06.425548

**Authors:** Ilan N. Rubin, Iaroslav Ispolatov, Michael Doebeli

**Affiliations:** Department of Zoology, University of British Columbia, Vancouver, British Columbia, Canada; Departamento de Física, Universidad de Santiago de Chile, Santiago, Chile; Department of Mathematics, University of British Columbia, Vancouver, British Columbia, Canada

## Abstract

One of the oldest and most persistent questions in ecology and evolution is whether natural communities tend to evolve toward saturation and maximal diversity. Robert MacArthur’s classical theory of niche packing and the theory of adaptive radiations both imply that populations will diversify and fully partition any available niche space. However, the saturation of natural populations is still very much an open area of debate and investigation. Additionally, recent evolutionary theory suggests the existence of alternative evolutionary stable states (ESSs), which implies that some stable communities may not be fully saturated. Using models with classical Lokta-Volterra ecological dynamics and three formulations of evolutionary dynamics (a model using adaptive dynamics, an individual-based model, and a partial differential equation model), we show that following an adaptive radiation, communities can often get stuck in low diversity states when limited by mutations of small phenotypic effect. These low diversity metastable states can also be maintained by limited resources and finite population sizes. When small mutations and finite populations are considered together, it is clear that despite the presence of higher-diversity stable states, natural populations are likely not fully saturating their environment and leaving potential niche space unfilled. Additionally, within-species variation can further reduce community diversity from levels predicted by models that assume species-level homogeneity.

**Author summary:** Understanding if and when communities evolve to saturate their local environments is imperative to our understanding of natural populations. Using computer simulations of classical evolutionary models, we study whether adaptive radiations tend to lead toward saturated communities, in which no new species can invade or remain trapped in alternative, lower diversity stable states. We show that with asymmetric competition and small effect mutations, evolutionary Red Queen dynamics can trap communities in low diversity metastable states. Moreover, limited resources not only reduces community population sizes, but also reduces community diversity, denying the formation of saturated communities and stabilizing low diversity, non-stationary evolutionary dynamics. Our results are directly relevant to the longstanding questions important to both ecological empiricists and theoreticians on the species packing and saturation of natural environments. Also, by showing the ease evolution can trap communities in low diversity metastable states, we demonstrate the potential harm in relying solely on ESSs to answer questions of biodiversity.

## Introduction

One of the fundamental goals of ecology is to understand how biodiversity is maintained. Competition theory predicts that two species with identical niches will lead to competitive exclusion and one will win out. Early theoretical work by Robert MacArthur [1–3] codified the idea that competition can lead to the partitioning of continuous phenotype space into niches, allowing for the stable coexistence of species. For species to stably coexist, selective pressures will limit similarity and partition species into individual niches.

MacArthur introduced the idea of niche packing as one of two ways of diversifying (the other being exploration) [1]. Niche packing implies that a higher density of species must lead to a greater partitioning of the available niche space. This has inevitably led ecologists to ask at what point communities will saturate with maximal diversity and whether natural communities tend to exist at saturation [4–6]. While this question has recently led to vigorous debate and research, there has been little theoretical treatment of the evolutionary dynamics for saturated versus unsaturated communities. For instance, it is a well known result that ecological stability does not necessarily imply evolutionary stability and that maximal ecological diversity is often evolutionarily unstable [7].

It has become increasingly clear that eco-evolutionary dynamics play a large role in the long-term maintenance of biodiversity. For example, adaptive radiations, the rapid ecological differentiation of a single clade [8], are able to generate vast amounts of diversity [9,10]. Eco-evolutionary models of frequency-dependent competition with mutation have been used to show how adaptive radiations can emerge from these simple competitive interactions, leading to diversification and niche partitioning [11,12].

Recent work has investigated the theoretical existence of alternative evolutionary stable states (ESSs – long-term endpoints of an evolutionary process), the presence of multiple different communities in a given system that are uninvadable by a mutant of small effect [13–15]. The presence of alternative ESSs necessarily implies that certain stable ecological communities may not be at saturation. Certain ESSs may even be “Garden of Edens” that are unreachable by successive small mutations [16] or are only reachable through rapid evolution that occurs on the same timescale as ecology [13,14]. Calcagno et al. [15] create an atypical scenario running contrary to classical niche-partitioning reasoning, where the initiation of diversification is dependent on there already being diversity present. While both of these results are intriguing, the simple question of whether adaptive radiations tend toward saturated communities or stall at an unsaturated ESS remains largely unanswered.

Given the extraordinary complexity of biological processes, it is natural to think that selection takes place in many dimensions. Despite this, a majority of our intuition of evolutionary dynamics come from narratives of individual traits or models with single phenotypic dynamics. Recent studies of evolution in high-dimensional phenotype space show that evolutionary dynamics can often be complex [17,18]. With increasing dimension, low and intermediate levels of diversity are increasingly non-stationary, with periodicity most common in lower dimensions and chaos in high dimensions. As the community diversifies, evolutionary dynamics slow down, often, but not always, fully stabilizing [18]. While the patterns that emerge are often stable, it is not yet clear whether these represent fully saturated communities or lower diversity ESSs. This question is essential for our understanding of how diversity is generated and maintained in natural communities.

Here we investigate whether different patterns of niche partitioning in multi-dimensional phenotype space can lead to alternate levels of stable diversity in a given system and whether adaptive radiations generally lead to saturated communities. We use an eco-evolutionary model with ecological dynamics described by the classic formulation of Lotka-Volterra competition and evolution as a trait substitution process in continuous phenotype space. For computational and visualization simplicity, only two-dimensional phenotypes are considered. The evolutionary dynamics are solved using three separate modeling frameworks: adaptive dynamics [19], individual-based simulations, and partial differential equations. All three models are run with the same ecological dynamics. Numerical simulations of the adaptive dynamics allow for the most efficient computation and most extensive exploration of the three modeling frameworks. The individual-based models allow us to test the assumptions of adaptive dynamics and explore the effects of finite population size and phenotype distributions on the patterns of adaptive radiations.

We will show that adaptive radiations in multiple dimensions often lead to locally stable levels of diversity. Higher diversity states and eventually globally stable community saturation may be reached either through large mutations, immigration of species with phenotypes novel to the community, or if the adaptive radiation was initiated with higher levels of standing genetic variation. With asymmetric competition, the lower diversity states often take the form of stable limit cycles that represent Red Queen dynamics [20]. While these low diversity states are only locally stable, finite population sizes further restrict diversification despite available niche space, stabilizing low diversity, unsaturated communities and, depending on the system, perpetuating Red Queen dynamics. These patterns of locally stable, low levels of diversity are likely ubiquitous in nature, leaving communities unsaturated and areas of phenotype space open for invasion and continued adaptation. These results shed light on the speed and characteristics of adaptive radiations as well as whether or not diversity tends to evolve to saturation [6].

## Models and methods

### Ecological Dynamics

Here we examine a model of phenotypic evolution based on classic logistic Lotka-Volterra ecological dynamics. Individuals are defined by a two-dimensional continuous phenotype and population size. In a monomorphic population of a single phenotype 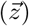, the equilibrium density of that population equals the carrying capacity, 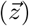, of that phenotype. We will use two different forms of carrying capacity functions, from now referenced to as quartic or radially symmetric (Fig 1).

**Fig 1.**
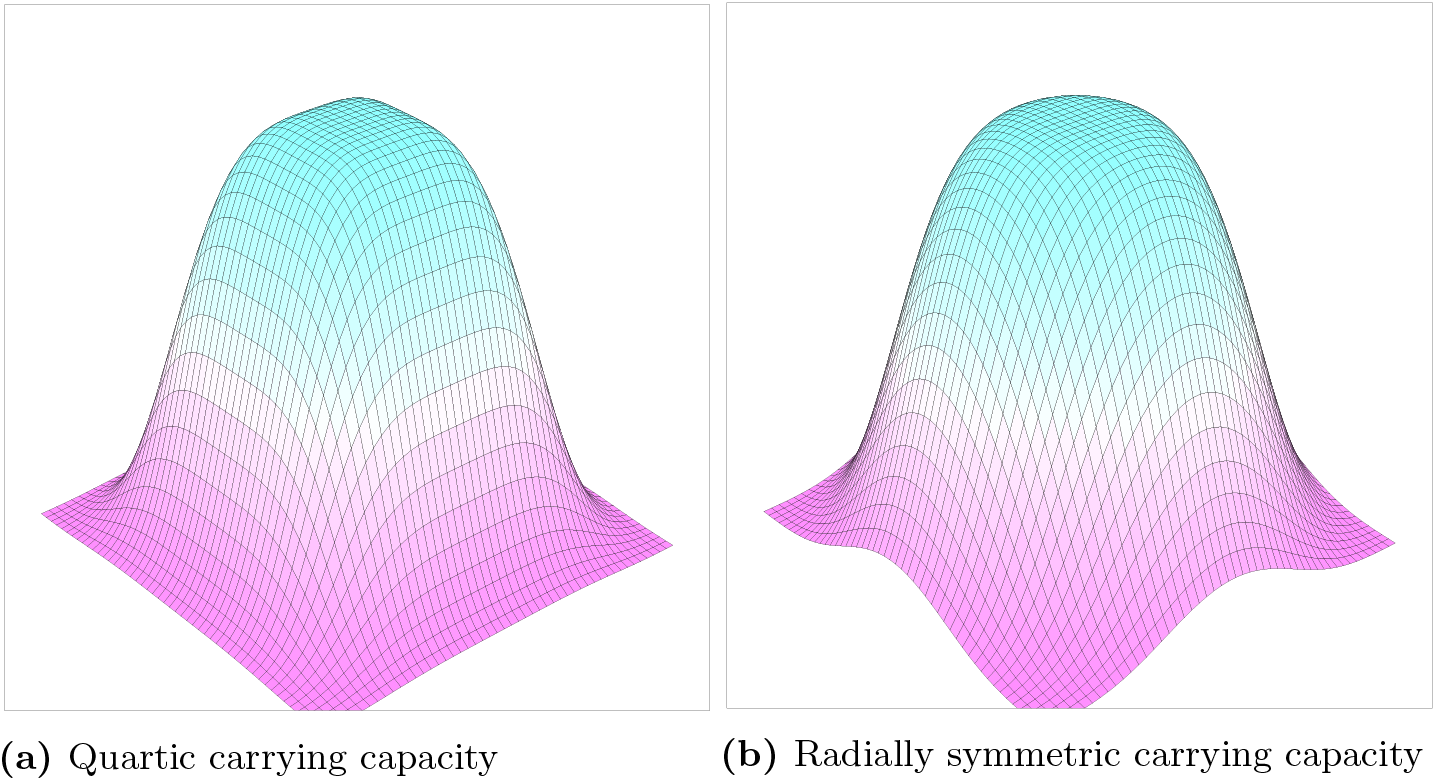
Carrying capacity functions. Two carrying capacity functions in 2D trait space. The peak of both functions is 1 at the origin and decreases to 0 as the phenotype increases or decreases. The quartic carrying capacity has a square peak while the radially symmetric carrying capacity has a circular peak. As both functions are of order 4, they are “flatter” on top than a standard Gaussian distribution. For the individual based simulations, the same carrying capacity functions are used, but multiplied by a scalar *K_max_* that determines carrying capacity in a number of individuals at the origin. This scalar controls the “richness” of the environment.

For the quartic case the carrying capacity function is

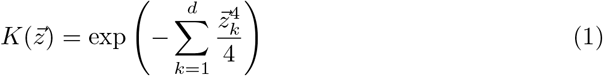

The radially symmetric carrying capacity is

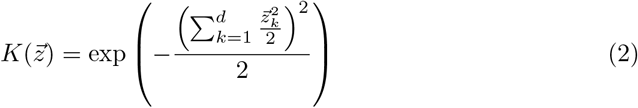

When evolution is considered, both these functions impose stabilizing selection on the phenotype towards the origin where the carrying capacity is maximal. Due to the higher order term in both of these functions, carrying capacity has a fairly “flat” peak, which naturally restricts viable phenotype space to approximately between −2 and 2 in each dimension. In addition to naturally limiting viable phenotype space, using a quartic carrying capacity avoids the structural instability of a Gaussian carrying capacity and Gaussian competition kernel that results in infinite branching [21, 22].

Individuals compete with others governed by a competition kernel 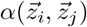. The competition kernel equals 1 when an individual competes with another with the same phenotype and decreases to 0 as the phenotypic similarity of the two competing individuals decreases. Thus, similar individuals will have a greater effect on each other’s growth compared to individuals with more distinct phenotypes. We consider situations with both symmetric and asymmetric competition. Symmetric competition refers to when individuals impart exactly the same competition load on each other such that 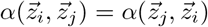. While this is traditional [21] and conceptually convenient, perfect symmetry rarely occurs in nature and asymmetric competition has been explicitly measured [23]. Therefore, we also consider asymmetric competition where 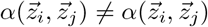. For symmetric competition, the competition kernel takes the form of a Gaussian with variance equals to 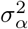. For asymmetric competition an term is added with coefficients *b* that determine the nature of the competitive interaction between the phenotypes in the two dimensions. This asymmetric term is non-mechanistic and can be thought of as the first-order term in a Taylor expansion of some higher order asymmetric interaction function [17]. In this way, it is the simplest way to add asymmetry to Gaussian competition.

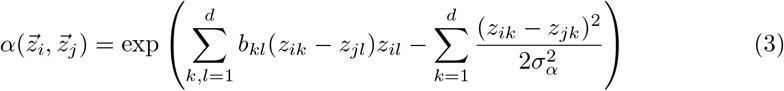

When the *b* coefficients are set equal to 0, the function reduces to symmetric, Gaussian competition. The competition kernel provides the frequency-dependent component of the ecological dynamics, which allows for the stable coexistence of multiple competing phenotypes under certain conditions.

While the functional forms used here for carrying capacity and competition are largely phenomenological, they represent biologically reasonable scenarios [21] and are well supported in the literature [21,24].

Based on the classic Lotka-Volterra formulation, ecological dynamics are thus:

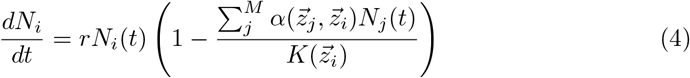

for growth rate *r*, a population *i* with phenotype 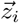 and population size *N_i_*(*t*) that competes with each of *M* other groups of individuals with distinct phenotypes 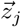, and population sizes *N_j_*(*t*).

For simplicity and computational tractability, all simulations are spatially homogeneous and positions of individuals or groups in space are ignored.

### Evolutionary Dynamics

#### Adaptive Dynamics

We model evolution using adaptive dynamics. Adaptive dynamics allows for tractable computation of evolutionary dynamics when a few basic assumptions are met: (1) there is a 1-to-1 map from genotype to phenotype; (2) all genetically identical individuals can be represented as a single phenotype with no phenotypic variation (i.e., a delta function); (3) when a favorable mutation arises, it out-competes the resident, driving the resident extinct; and (4) mutations are small and rare. It is essential in adaptive dynamics derivation that the resident populations are at their ecological equilibrium, or in other words, that population dynamics are infinitely fast on the evolutionary timescale and any mutant that arises either fixes in the population or goes extinct before another mutant is introduced.

To derive the adaptive dynamics, we must first define the invasion fitness [25] 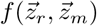 as the per capita birth rate of a rare mutant *m* in the monomorphic population of resident *r* that is at its ecological equilibrium population size 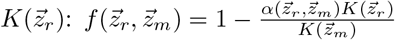. When the invasion fitness is positive 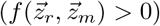, the mutant can invade the resident population. When mutual invasibility occurs, i.e., the invasion fitness of the resident into an equilibrium population of the mutant is also positive 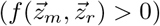, individuals of the two phenotypes can coexist indefinitely.

By taking the partial derivative of the invasion fitness function with respect to the mutant when the mutant phenotype equals the resident (mutations are infinitely small) we can derive the selection gradient and then the adaptive dynamics, which describe the trajectory of a single, monomorphic population as it evolves in the trait space (which in our case in two-dimensional). A more detailed description of adaptive dynamics, please refer to Dieckmann and Law [26] and Doebeli [21].

#### Speciation

The canonical equation of adaptive dynamics, as described above, can thus only describe the movement of populations in phenotype space, but not speciation or extinction events. Evolutionary branching is a well known phenomenon that occurs when there is an attracting equilibrium or nullcline [27–29] in trait space that is also a fitness minimum [19]. To model these we use a well described algorithm [18] where the adaptive dynamics are run for some period of time at which a random population is chosen and a small mutant is introduced nearby. If that mutant is viable and is mutually invasible with its parent, the population successfully branched and the simulation continues with an additional phenotype. In doing so, this algorithm results in deterministic ecological dynamics and quasi-deterministic evolutionary dynamics in which evolutionary trajectories are deterministic but branching is stochastic.

#### Simulations

Evolutionary simulations are conducted thusly: (1) solve for the ecological equilibrium of the current population; (2) delete any populations that fall below a minimum viable population size and considered extinct; (3) solve the adaptive dynamics for a fixed length of evolutionary time; (4) for computational speed, merge any two populations whose phenotypes are within a very small distance of each other (Δ*z*); (5) introduce a mutant population, whose phenotype is a small deviation (*ε_mut_*) from the parent in a random direction; (6) delete the mutant if it cannot invade the population; and (7) repeat the process until an evolutionary stable community or a designated time has been reached. After each time step, similar phenotypes (those with phenotypes within a small distance from each other) are clustered into species as to better represent the number of distinct phenotypic groups alive at any given time (this does not affect the simulations and is only an accounting device).

### Stability

Given the mathematical complexity of these dynamics, we were unable to analytically determine the evolutionary stability of the resulting communities. Instead, we define metastable evolutionary states, stationary or cyclic, as those, to which the system quickly converges and in which it resides much longer than the convergence time, possibly indefinitely. Mechanistically, the exit from a metastable state is conditional on significantly larger mutations than the convergence to it.

Our definition of metastability is in line with more common causes in physics and chemistry. Consider for example a well-studied problem of protein folding [30]: A model initiated with a completely disordered state quickly folds into one of potentially very many metastable conformations, and it takes long time or a potential guidance (assistance) from other proteins (chaperons) to reach the native functional fold. However, there is a subtle difference between metastability in ordering in physics and metastable states in evolution. In the physics systems, there usually exists a function, such as a free energy, that reaches its absolute minimum in the truly stable state, while the metastable states correspond to local minima, providing a very clear distinction between the former and the latter states. However, in essentially non-stationary systems, such as evolving systems, it is often impossible to define such a function.

### Individual-Based Model

In order to examine how finite population size affects the evolutionary dynamics, we use an individual-based simulation. In these simulations, individuals have a fixed birth rate and a frequency-dependent death rate calculated from the carrying capacity and competition kernel as described above. Every time step a single individual is chosen to die or give birth to an offspring with phenotype equal to its parent plus a small mutation in a random phenotypic direction. Instead of a single phenotype describing each population of individuals, populations are now represented by a cloud of points in phenotype space. In order to determine distinct species, individuals are clustered such that every individual of a given “species” lies within a prescribed phenotypic distance to at least one other member of that species. This clustering is purely accounting and has no bearing on the dynamics. For a more detailed account of individual-based simulations and their relation to adaptive dynamics please see Champagnat et al. [31, 32] and Ispolatov et al. [33].

The individual-based simulations are fully stochastic and do not require any of assumptions of adaptive dynamics except the 1-to-1 genotype-phenotype map. Thus, in addition to providing a means to investigate the effects of finite population size, these simulations also act as a check on the applicability of the assumptions required by adaptive dynamics. Additionally, unlike adaptive dynamics, evolutionary branching is an emergent property of the individual-based simulations [31] and acts as a further confirmation of diversification within the adaptive dynamics simulations.

For both adaptive dynamics and individual-based simulations the population is initially seeded with individuals with a randomly chosen phenotype between −2 and 2 in each dimension, roughly coinciding with the area of viable phenotype space for both carrying capacity functions. Simulations were seeded with different numbers of individuals, each with a distinct phenotype, in order to investigate the effects of standing genetic variation on the dynamics of adaptive radiation.

From here on we will refer to groups of phenotypically similar individuals as a species. In the individual-based model a species takes the form of a distinct cloud of points while in the adaptive dynamics simulations a species is represented by a single phenotype and its population size.

## Results

Because the carrying capacity functions we investigate here both have a broader maximum at the origin than that of the Gaussian competition kernel, the origin is always the minimum of the invasion fitness and there necessarily exists a branching point at the origin. Thus the possibility of evolutionary coexistence between two or more phenotypes is guaranteed. In order to further encourage diversification, we only use *σ_α_* < 1, which would be the condition to generate a branching point at the origin if we had instead used carrying capacity functions of equal order to the competition kernel.

Unless otherwise noted, all figures were generated using parameters listed in the Supplementary Information, S1 Appendix.

### Symmetric competition

For symmetric competition (*b* = 0), a quartic carrying capacity, and *σ_alpha_* = 0.5, when starting with a single species (monomorphic population defined by its two dimensional phenotype), that species quickly evolves under directional selection toward the fitness maximum at the origin. Once there, it will start to successively branch, quickly stabilizing into an evolutionary stable community (ESC) with 16 species arrayed in a 4×4 grid (Fig 2). This pattern naturally emerges as the community evolves to pack the viable (and approximately square) niche space, while still maintaining space from neighboring species due to competition. For a more detailed discussion of the patterns and dynamics of species packing in multi-dimensional phenotype spaces please refer to [18].

**Fig 2.**
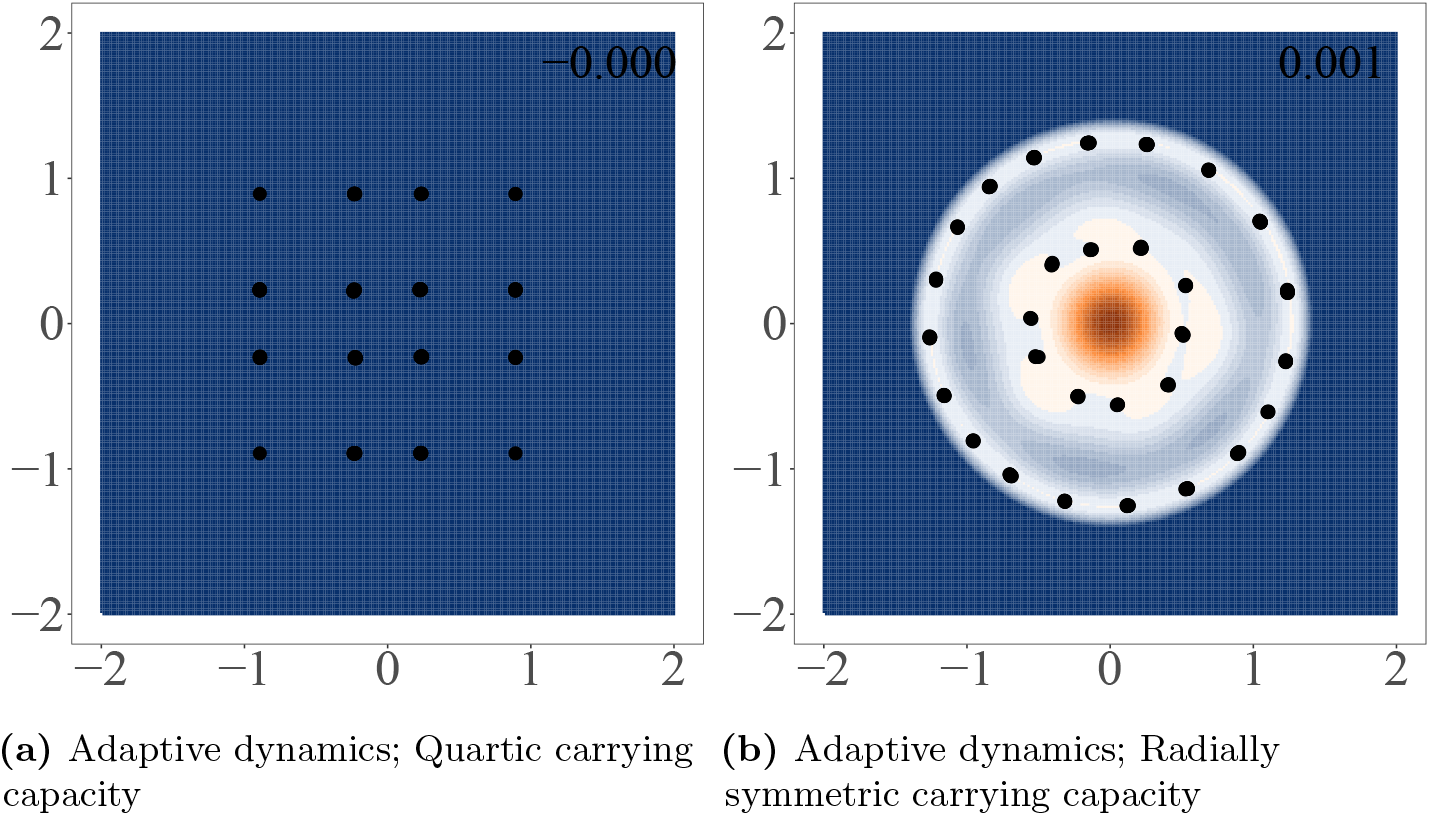
Stable states for symmetric competition with quartic and radially symmetric carrying capacity. Figures were generated using adaptive dynamics simulations. Points in the upper panels represent surviving species at the end of each simulation and the surface shows the invasion fitness (per capita growth rate of a rare mutant) with positive invasion fitness displayed in orange and negative in blue. The maximum invasion fitness for each panel is printed in the top right corner. The range displayed for all axes is −2 to 2. When a symmetric competition kernel was used, simulations all converged on similar patterns regardless of the initial population. All simulations are run with the same parameters, which can be found in the Supplementary Information.

With a radially symmetric carrying capacity, the story is largely the same, but instead of diversifying into a grid, the population arranges itself into two concentric circles. The invasion fitness along the ridges of these concentric circles is nearly flat and equal to 0. This resulted in nearly neutral selection along the ridges and different numbers of final species in each simulation, ranging from 24 to 34 between both circles (Fig 2). While the invasion fitness landscape in figure also indicates directional selection toward the origin, the invasion fitness at the origin is three orders of magnitude less than maximum invasion fitness during the initial adaptive radiation, indicating weak selection. Manually placing a population at or near the origin does result in a stable configuration, but diversification toward the origin is so slow it was nearly imperceptible during our simulations even when simulations were run for far longer than our results shown here. Like selection toward the origin, areas of slight positive invasion fitness on the concentric circles indicate that these configurations are likely not fully stable. Clusters on the outer ring seem to be slowly continuing to undergo diversification as long as the simulations were run, though the rate of speciation events and speed of evolution slowed dramatically as the rings filled up.

These simulations with the radially symmetric carrying capacity function and symmetric competition were the only simulations we ran that did not fully settle in stationary or cyclic configurations on timescales that were computationally feasible. Similar long-term transients were not found in any simulations with different functional forms for carrying capacity or competition, or with different parameterizations, so it is likely that this is a degenerate case caused by a perfectly radially symmetric carrying capacity function and Gaussian competition. Indeed, it is a well known result that in a one-dimensional trait space with a Gaussian carrying capacity kernel and symmetric, Gaussian competition, adaptive dynamics results in an infinite branching process [21]. However, like in our simulations, this “infinite branching” quickly degrades to discrete phenotypes when either the carrying capacity or competition kernels are altered from being perfectly symmetric.

### Asymmetric competition can lead to Red Queen dynamics

For certain values of the four *b* coefficients (the coefficients that govern the nature of asymmetric competition) the population quickly settles into an evolutionary stable state (ESS). As mentioned in Doebeli et al. [18], in a two-dimensional system like the one simulated here, most randomly chosen b values result in evolutionarily stable communities (ESCs). These configurations are often grid-like for the quartic carrying capacity or concentric circles for the radially symmetric carrying capacity but with some “skew” related to the asymmetry in competition. These ESCs can fully saturate the environment, leaving little to no area of trait space with positive invasion fitness (Fig 2).

Other combinations of b however, can result in non-equilibrium evolutionary dynamics. There is significant literature already detailing how asymmetric competition can lead to non-equilibrium dynamics – either stable limit cycles [34] or in higher dimensions, chaos [17,18,35]. These stable limit cycles represent Red Queen dynamics [34] where the community of one or more species continuously evolves (Fig 3). Notably, these cycles are not driven cyclic dynamics in population size, which are not possible with purely competitive interactions and logistic growth, but by the asymmetric competition driving selection around the periodic orbit as can be seen by the selection gradient in figure 3.

**Fig 3.**
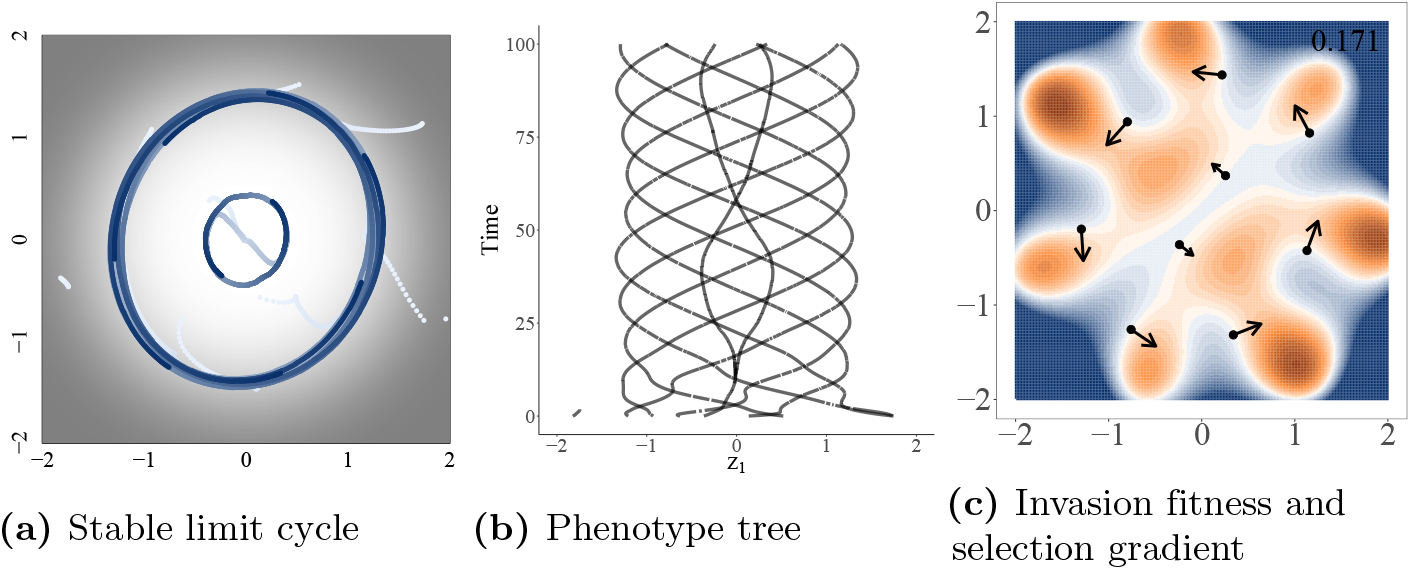
Asymmetric competition can lead to Red Queen dynamics. Red queen dynamics denotes a situation in which one or more populations continuously evolve on a a stable limit cycle in phenotype space. Simulation were run using the radially symmetric carrying capacity and initiated with 10 random species. Panels A and B show the complete history of evolutionary dynamics, with time in panel A increasing from white to blue and carrying capacity increasing from black = 0 to white = 1. Panel C is a depiction of the population at the end of the simulation. Colors in panel C represent the invasion fitness (per capita growth rate of a new mutant if it were to arise). Positive invasion fitness is shown in shades of orange (maximum of 0.17), negative in blue, and invasion fitness equal to zero in white. Arrows are proportional to the square root of the selection gradient for each species. Simulation time was cut to only 100 time steps in the figure so the limit cycles could be more easily seen.

### Red Queen dynamics can lead to alternate levels of metastable diversity

Notably, in simulations that result in stable limit cycles, different numbers of species may emerge (Fig 4). This emergent diversity is then maintained. Unlike an ESC, these stable limit cycles have large areas of positive invasion fitness that are reachable by small mutations – a fact that underlies the non-zero selection gradient and drives the cyclic movement. This also means that mutants can continuously invade. However, all areas of positive invasion fitness reachable by small mutations only perpetuate the oscillatory motion rather than initiating diversification.

**Fig 4.**
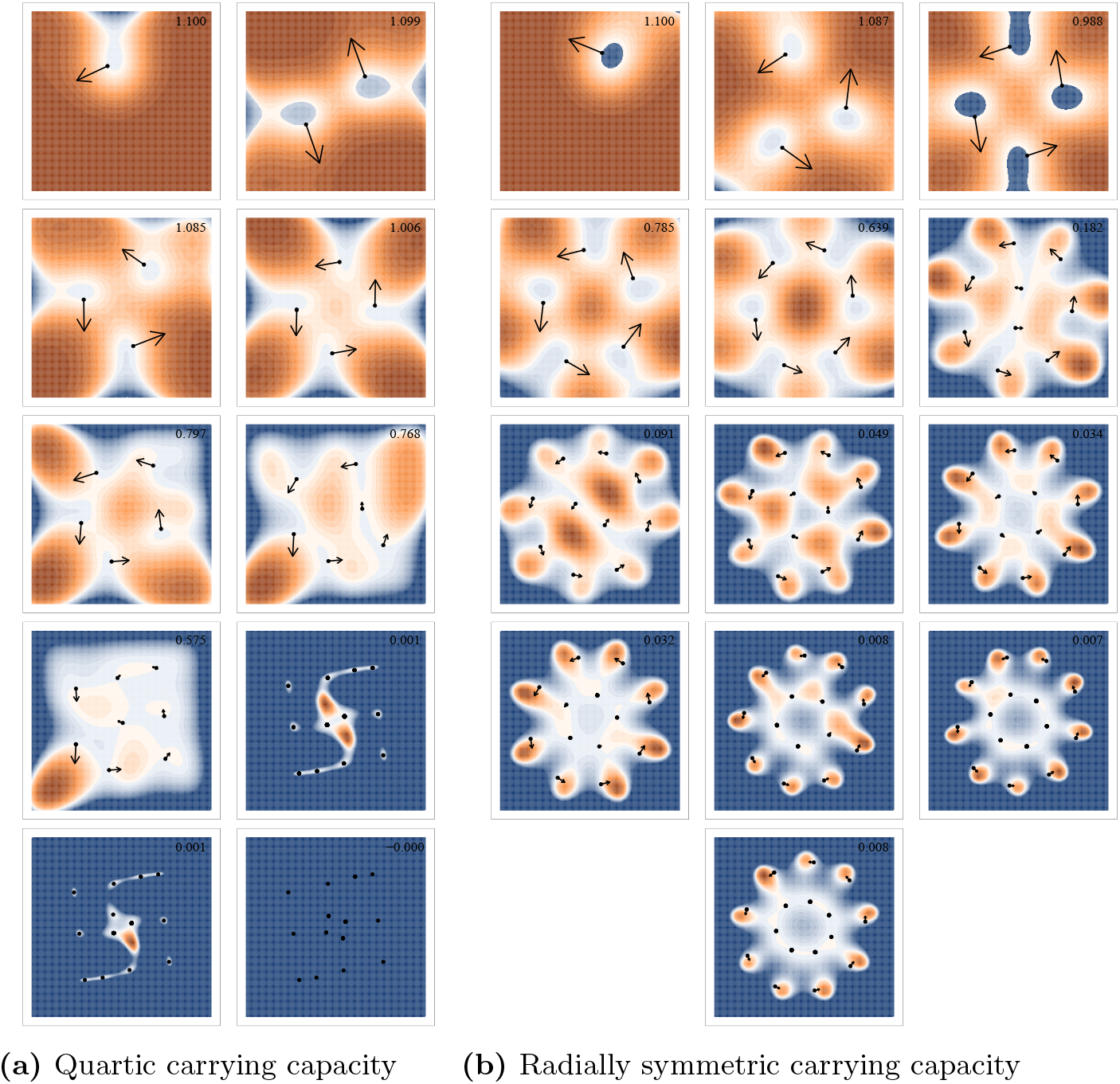
Invasion fitness landscapes for alternative metastable states. Alternative metastable states resulting from simulations with different levels of initial population diversity. Each panel represents an alternative level of metastable diversity for a single set of parameters and only differ based on the randomly generated initial communities. Points represent surviving species at the end of each simulation. Arrows are proportional to the square root of the selection gradient for each species. The surface shows the invasion fitness (per capita growth rate of a rare mutant) with positive invasion fitness displayed in orange and negative in blue. The maximum invasion fitness for each panel is displayed in the top right corner of the panel. All axes are displayed from −2 to 2. Simulations in the left two columns use the quartic carrying capacity and those in the right three columns use the radially symmetric carrying capacity. All other parameters can be found in the Supplementary Information. Additional versions of these figures with the initial population also display in addition to the final configuration are linked to in the Supplementary Information.

To examine this further, we seeded the adaptive dynamics simulations with an arbitrary number of randomly chosen phenotypes (between 1 and 100 initial species). When seeded with a high number of species, most die out immediately, but many survive. Even after the system is simulated for a long period of time (100,000 branching mutations attempted), those simulations seeded with few species maintain the low diversity while simulations seeded with more tend toward higher diversity stable limit cycles or ESCs (Fig 5). For the parameterization illustrated here and a quartic carrying capacity, metastable limit cycles with 1-6, or 8 species all emerged depending on the initial diversity in the simulation (Figs 4, 5). Evolutionary metastable states of 12, 13, or 14 clusters also emerged when seeded with high initial diversity. Other parameterizations of asymmetric competition (values for *b*) were also tested and while some parameterizations did not result in any cyclic dynamics, alternative levels of metastable diversity were always found. Interestingly, using the quartic carrying capacity, we were unable to find a parameterization of *b* that resulted in a stable limit cycle at low diversity but did not eventually saturate to an ESC if seeded with a diverse initial community.

**Fig 5.**
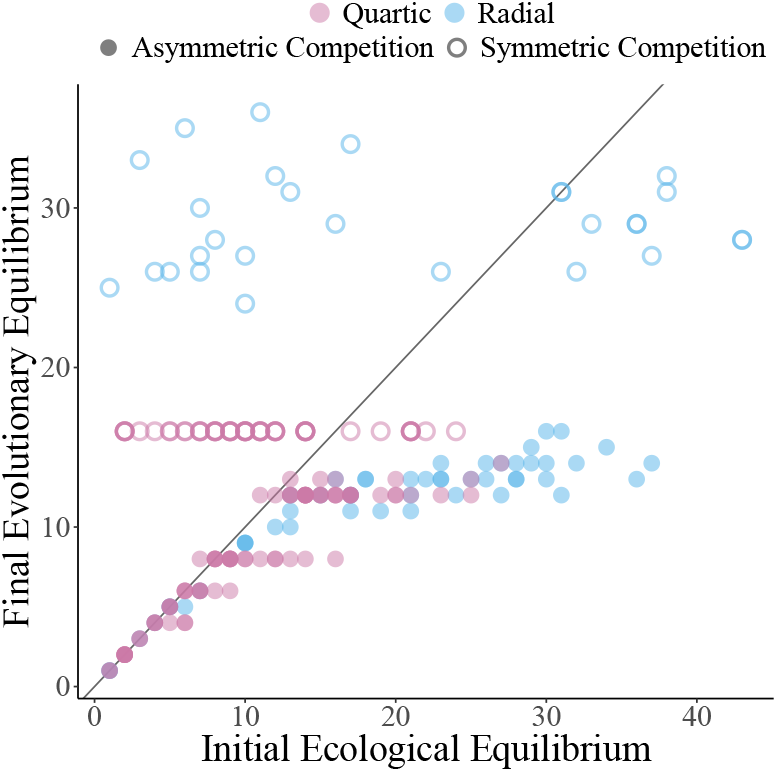
Levels of metastable diversity. The final evolutionary diversity when seeding the simulation with different numbers of initial species. Red indicates simulations with a quartic carrying capacity kernel, while blue are those with a radially symmetric carrying capacity kernel. Open circles are simulations with only symmetric competition. Full circles are simulations run with asymmetric competition. The b values dictating the competition asymmetry can be found in the Supplementary Information (S5A Table). All other parameters remained the same for all simulations and can be found in the Supplementary Information as well.

The presence of a high diversity ESS was not always the case with the radially symmetric carrying capacity. The pattern of low seeded diversity leading to low diversity metastable oscillations is the same, but high diversity stable configurations were often oscillatory as well. Many different simulations with randomly chosen asymmetric competition parameters and the radially symmetric carrying capacity seemingly had no fully saturated ESC, including the parameterization shown here (Fig 4).

### Large mutations pushes populations from lower diversity cycles toward saturated ESS

The small mutation assumption of adaptive dynamics is inflexible and necessary in order to derive the evolutionary dynamics. However, branching events are simulated manually. Therefore, we can increase the size of branching mutations (instead of a small fixed mutation *ε_mut_*, mutant phenotypes are chosen from a Gaussian distribution with mean equal to the parent and standard deviation *σ_mut_* ≥ *ε_mut_*).

With a quartic carrying capacity, when *σ_mut_* is increased to 0.05 or larger the lower diversity metastable limit cycles eventually break down and the species-packed ESS is reached (Fig 6). This indicates that these lower diversity cycles are locally stable while the high density ESS is globally stable. For the asymmetric parameters chosen here, the saturated ESS contains 14 species (Fig 4). Of note, because branching mutations are modeled as a Gaussian distribution, even with a small *σ_mut_* transitions from lower to higher diversity limit cycles did occur, but exceedingly rarely, allowing the lower diversity meta-stable states to persist until the end of our simulations.

**Fig 6.**
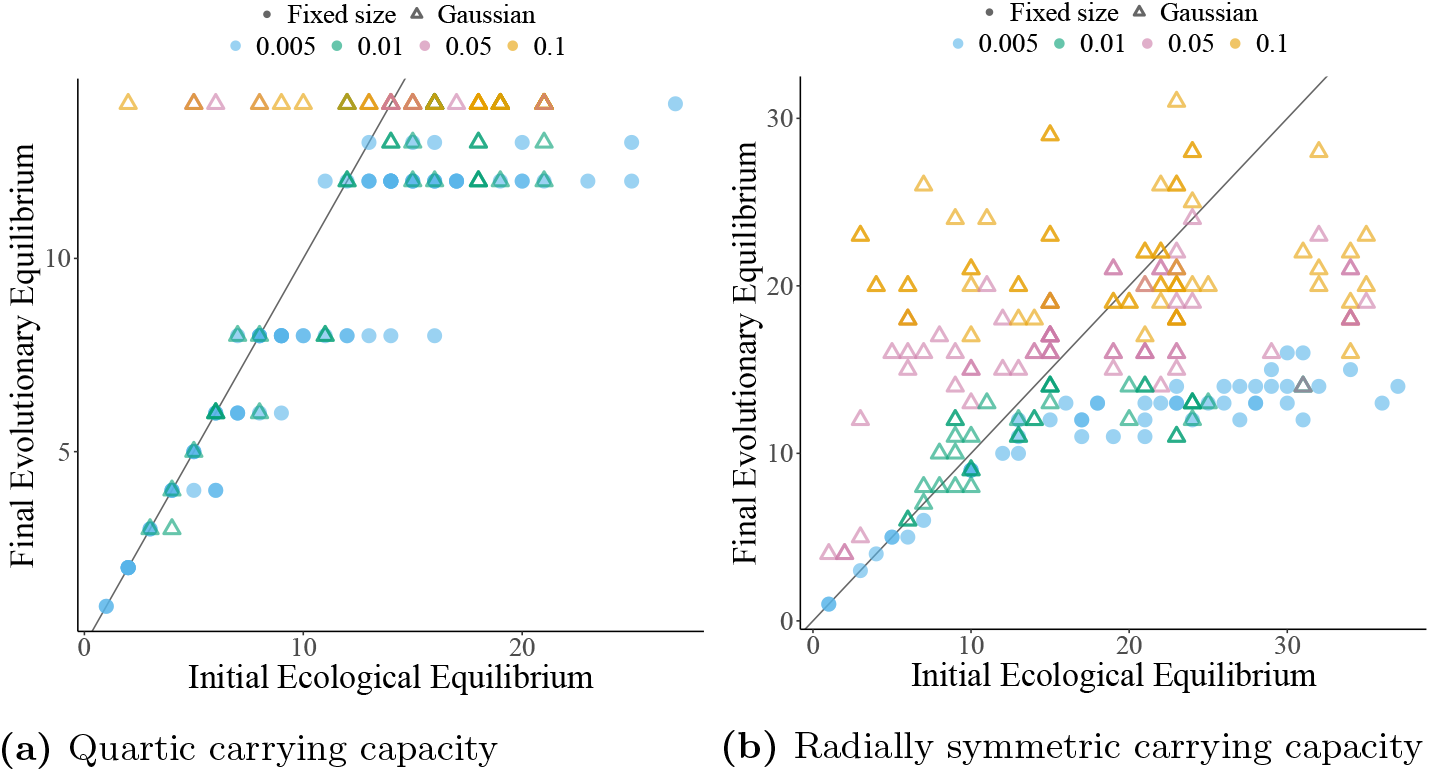
Large mutations allow escape from low diversity meta-stable states. The final evolutionary diversity when seeding the simulation with different numbers of initial species for differently sized mutations and asymmetric competition. For solid points, mutants were placed a small fixed distance away from the parents. For hollow triangles, mutations are drawn from a Gaussian with mean equal to the parent’s phenotype and standard deviation indicated by point color. Mutation size or standard deviation (depending on the mutation algorithm) are represented by color. All other parameters are the same as those listed in the Supplementary Information.

For radially symmetric carrying capacity simulations without an ESS, each parameter combination seemingly has a globally stable limit cycle that is eventually reached with large enough mutations. For the parameter values used as an example in this paper, this globally stable limit cycle has 8 species on the outer ring and 5 on the inner, both which cycle clockwise. Unlike with the quartic carrying capacity, there also exist metastable limit cycles with higher diversity than the globally stable one. These “super saturated” communities also collapse to the globally stable cycle when mutation size is increased. Super saturated stable limit cycles occurred rarely when seeded with a random initial species, but were easy to manufacture by manually placing species in trait space on approximately the two concentric circles that emerge in any community with more than six distinct species (Fig 4).

Because the globally stable community undergoes non-equilibrium dynamics, there still exist areas of positive invasion fitness that drive these cycles (Fig 7). Mutants into these areas of positive invasion fitness eventually out compete the nearby resident, returning the system to 13 species. The invasion fitness landscape around the inner ring is nearly flat, with shallow peaks, leading to nearly neutral local dynamics. This leads to the persistence of small clusters of mutants around the 5 inner species for relatively long periods of time (for a discussion on neutral coexistence see [36]). Despite this apparent increased diversity, the pattern of approximately 13 species clusters is maintained long-term.

**Fig 7.**
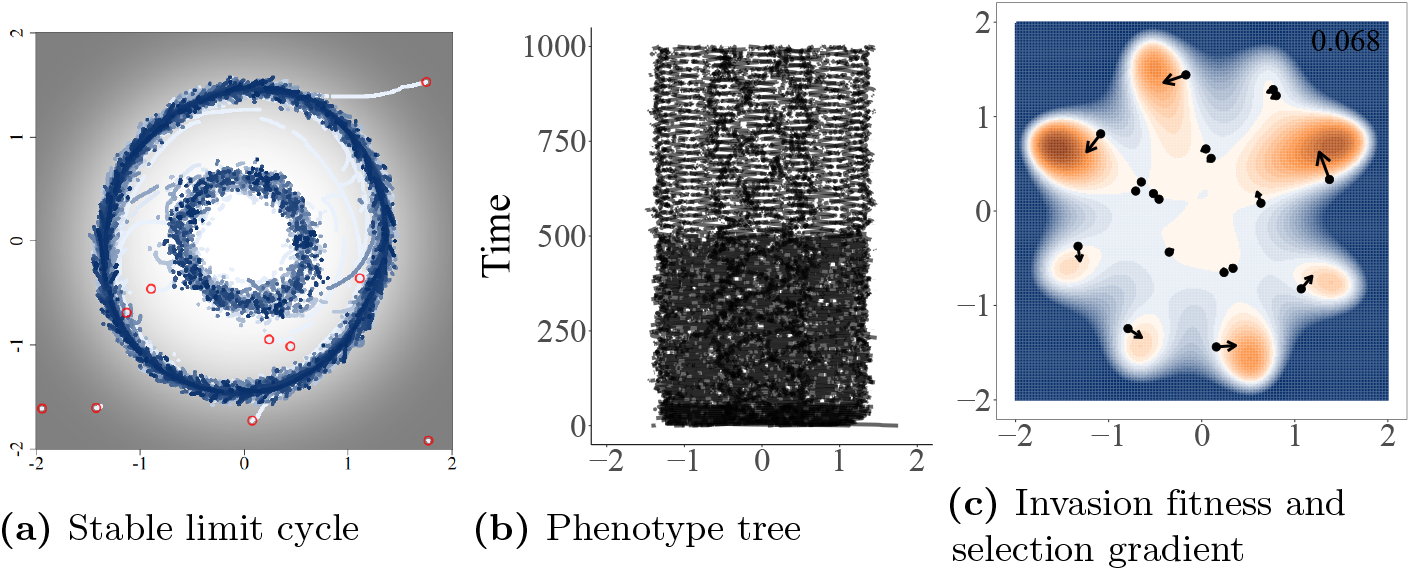
Large mutations can cause transitions between locally stable levels of diversity. Simulation were run using the radially symmetric carrying capacity, Gaussian distributed mutations with *σ_mut_* = 0.1, and 10 initial species (randomly chosen). Panels A and B show the complete history of evolutionary dynamics, with time in panel A increasing from white to blue and carrying capacity increasing from black = 0 to white = 1. The initial population is highlighted in red. Transitions between diversity states due to rare, large mutations can be seen in the change in frequency of the limit cycles in Panel B. Panel C is a depiction of the population at the end of the simulation. Colors in panel C represent the invasion fitness. Positive invasion fitness is shown in shades of orange (with a maximum of 0.068), negative in blue, and invasion fitness equal to zero in white. Arrows are proportional to the square root of the selection gradient for each species. Dynamics of the inner circle are under weak selection and provide the environment for mutants to persist for relatively long periods of time.

### Finite population sizes reduces maximal diversity and facilitates cycling

Individual-based simulations largely aligned with all the results of the adaptive dynamics (Fig 8). Because of the fully stochastic nature of the individual-based model, trajectories obviously didn’t match the adaptive dynamics exactly, but the numbers of phenotypic clusters and direction of oscillations in those simulations with limit-cycles qualitatively aligned as expected.

**Fig 8.**
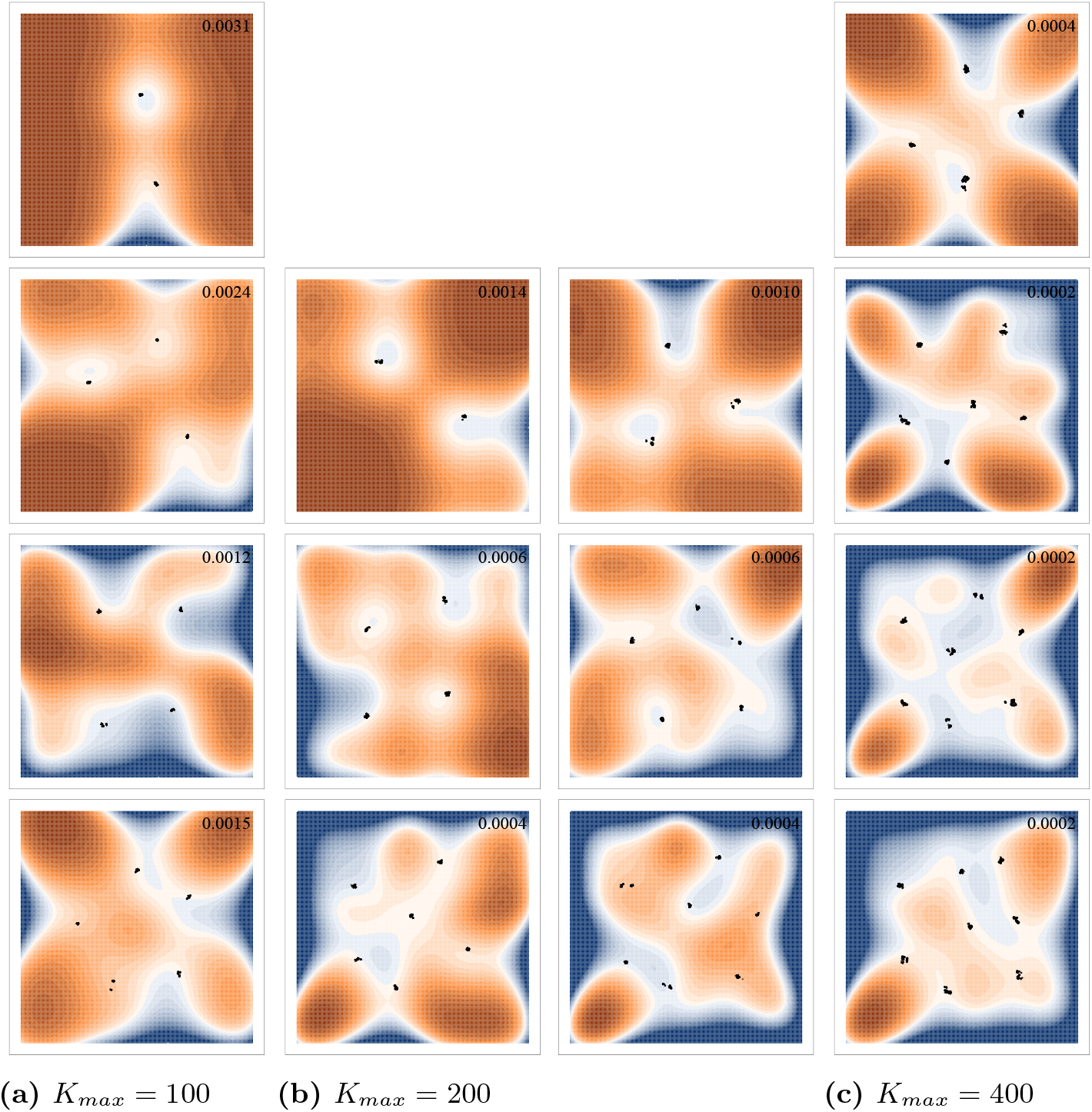
Alternative metastable states for different levels of final diversity in individual-based simulations. Points represent individuals at the end of each simulation. The surface shows the invasion fitness (per capita growth rate of a rare mutant) with positive invasion fitness displayed in orange and negative in blue. The maximum invasion fitness for each panel is displayed in the top right corner of the panel. All axes are displayed from −2 to 2. Simulations use a quartic carrying capacity with *K_max_* = 100 in the left column, *K_max_* = 200 in the middle columns, and *K_max_* = 400 in the right. All other parameters can be found in the Supplementary Information.

In these simulations, the overall size of the population can be regulated by a parameter *K_max_* that controls the height of the peak of the carrying capacity function at the origin. This does not set an artificial cap on the population and instead can be thought of as a parameter controlling the richness of the environment. When *K_max_* is large, there are sufficient resources available for a large population to grow. When it is small, resources are exhausted quickly and death rate increases, limiting the size of the population.

When we reduced *K_max_*, as expected the total size of the population decreased. More interestingly, the maximum number of phenotypic clusters that emerged decreased as well. For example, with a radially symmetric carrying capacity and asymmetric competition, when *Kmax* was set to 400, we see 7 or 8 species emerge when starting with high initial diversity (Fig 9). However, when *K_max_* = 200, no more than 6 species were ever maintained for a significant period of time. This was not due to demographic stochasticity, as those 6 species were often maintained in their limit cycle for very long periods of time without extinction or branching. This means that finite resources, and thus finite population sizes, limit the maximal diversity in a system, often facilitating cycling despite the presence of a theoretical global ESS.

**Fig 9.**
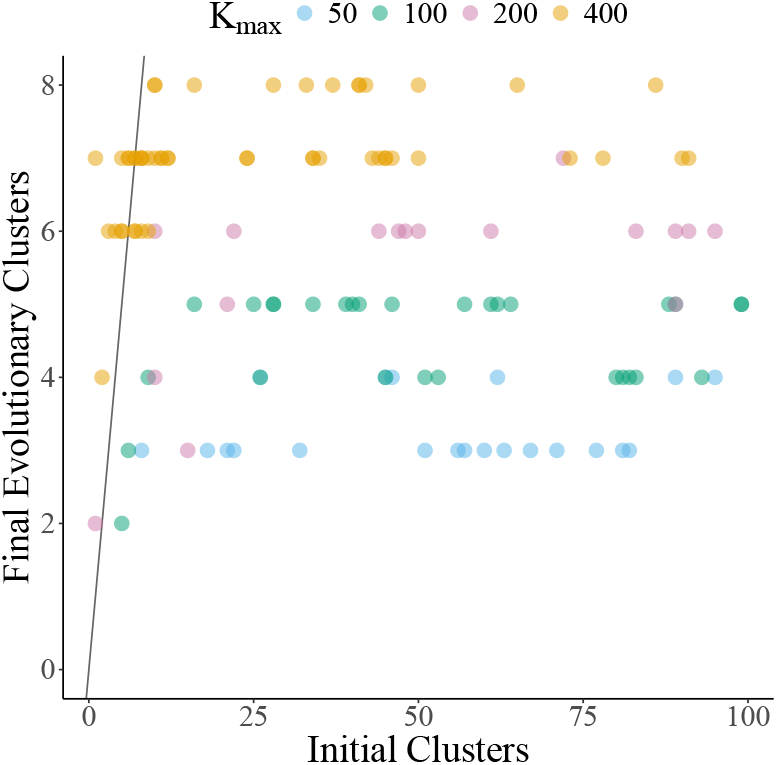
Finite population reduces realized diversity. The final number of phenotypic clusters for the individual based model when seeded with different initial population sizes. Individuals were clustered into groups by phenotypic similarity. To control for stochasticity, the final number of clusters was calculated as the median number of clusters over the last 200 time steps. Color indicates the maximum of the carrying capacity kernel (set at the origin). This represents the “richness” of the environment, with larger values modeling an environment with resources that are able to support a larger population. Simulations were run with a quartic carrying capacity kernel and all other parameters remained the same as previous simulations.

Despite the stochasticity, the same general pattern of locally stable, low diversity limit cycles with low initial phenotypic variation and high diversity limit cycles with high initial diversity largely persists (Fig 9). However, with a very low population size (e.g., *K_max_* = 50, Fig 8), demographic stochasticity overwhelms selection maintaining the low diversity limit cycles, allowing the population to consistently transition to a more stable, higher diversity state. Additionally, unlike in adaptive dynamics simulations, transitions from higher to lower diversity were also possible due to stochastic extinction (e.g., Fig. 10).

**Fig 10.**
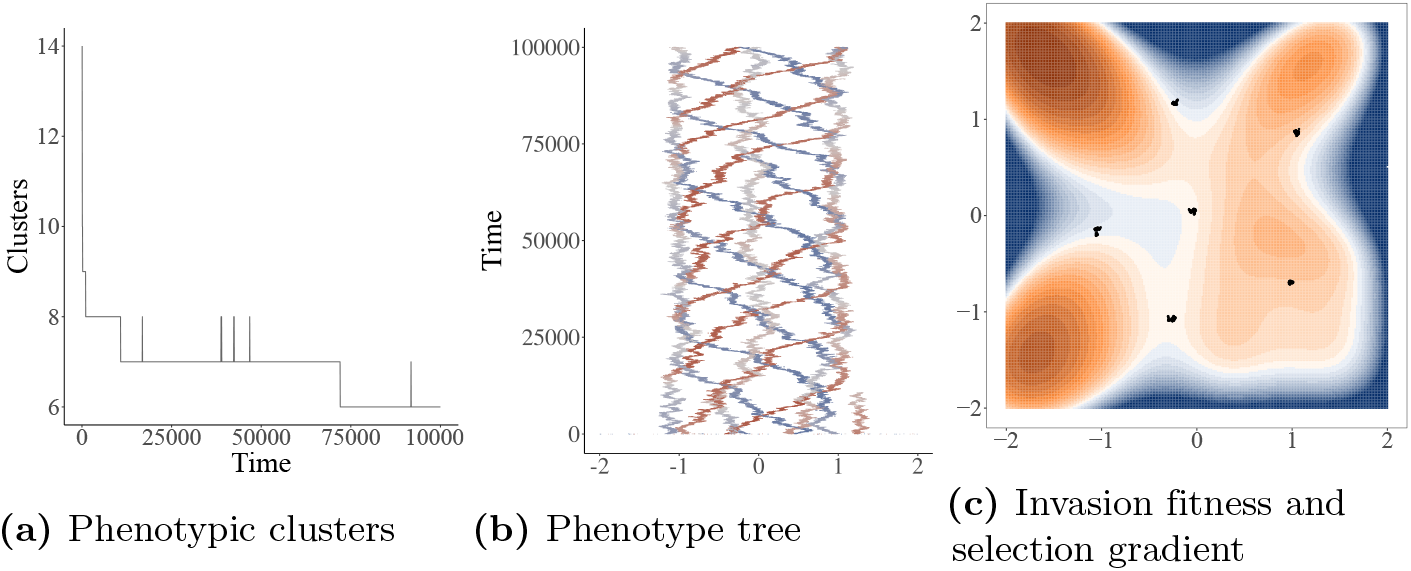
Finite population size can cause transitions in level of diversity due to demographic stochasticity. This simulation was run using the quartic carrying capacity, Gaussian mutation with *σ_mut_* = 0.005, *K_max_* = 200, and initiated with 89 randomly placed individuals. Panel A displays the number of phenotypic clusters over time. Panel B shows the complete history of evolutionary dynamics, with the x-axis representing phenotype *z*_1_ and color representing phenotype *z*_2_ (red = −2, white = 0, blue = 2). A transition between diversity states due to a stochastic extinction event can be seen approximately around time = 70000. Panel C is a depiction of the population at the end of the simulation (individuals shown as points). Colors in panel C represent the invasion fitness (measured as the birth rate - death rate of a new mutant). Positive invasion fitness is shown in shades of orange, negative in blue, and invasion fitness equal to zero in white.

### Small population sizes counteracts the effects of large mutations, maintaining low diversity limit cycles

With sufficiently large population sizes (*K_max_* ⪆ 400), increasing the size of mutations moderately had a similar effect to the adaptive dynamics simulations, allowing species to escape lower diversity limit cycles. However, larger mutations also increase the variation within species clusters. When mutation sizes were increased too much, distinct phenotypic clusters all but disappeared with individuals spread out across the entirety of the trait space. Even in these situations the population would cycle in the direction expected if a limit cycle exists.

In the adaptive dynamics, when mutations sizes increased, populations always converged on the most stable configuration, whether that be an ESS or a globally stable limit cycle. With small population size, even with increased mutation sizes, diversity levels remained low. This is because despite there being larger mutations allowing mutants to “jump” across areas of the negative invasion fitness and diversify, these new mutant species are rarely able to establish. The nature of finite resources inhibiting diversification is strong enough to counteract the ability of large mutations to escape locally stable limit cycles. In these cases, while the number of species was generally maintained, there was increased demographic stochasticity, with populations diversifying and going extinct more often than with smaller mutations.

## Discussion

We have shown here that systems based on Lotka-Volterra competition can cause many different levels of locally stable diversity emerge. This is particularly true with systems of asymmetric competition that lead to periodic evolutionary dynamics in phenotype space. These types of systems often get stuck in locally stable, low diversity limit cycles, despite the presence of a higher diversity global ESS or stable limit cycle.

Classic theory of adaptive radiations expects a quick burst of diversification, followed by a slowdown and possible settling to an ESS – a pattern that has also been shown in natural populations [8, 10, 37, 38]. Adaptive dynamics models show these exact dynamics with successive branching until an ESS is reached [21]. This ESS does not, however, imply community saturation. As expected, our results show that in an adaptive radiation, the population will quickly diversify. If a locally stable ESS or limit cycle is reached, diversification will then come to a stop. However, with small Gaussian mutations, eventually a rare mutation may be introduced that is able to invade the population and another diversification event takes place. This latest branching event could also trigger others, until a new locally stable community is reached. This means that during the early stages of an adaptive radiation, evolution is driven by relatively quick, successive, and small mutations, leading to the expected “early burst” of diversification. However, once a locally stable state is reached, rare, large mutations are necessary to “jump” areas of negative invasion fitness and initiate further diversification, followed by another round of successive small-effect mutations. This large effect mutation mirrors the classic idea of a “key innovation” [39], opening up new areas of adaptive opportunity.

Our findings also compliment findings of non-equilibrium dynamics in higher dimensions [17,18,35,40]. By manually restraining the levels of diversity, Doebeli et al. [18] were able to show that oscillatory and, in higher dimensions, chaotic dynamics are far more likely with lower levels of diversity. Here, we are able to generate the same non-equilibrium dynamics in two dimensions, but as an emergent property of the system. In our results, it became clear that while most systems had locally stable levelsof diversity, those with non-equilibrium dynamics were particularly difficult to escape. This is because in non-equilibrium systems, mutations into areas of positive invasion fitness tend to only perpetuate the same cycle. Instead of just having to jump canyons of negative invasion fitness in order to diversify, mutants likely also have to jump across a peak or saddle point of positive invasion fitness. Given the complexity of natural systems, evolution likely takes place in high dimensions. Taken together, these two results imply that many competitive ecosystems are likely unsaturated and undergoing some form of non-equilibrium, or Red Queen, dynamics. Indeed, previous results, both theoretical [41–43] and empirical [44], have implied that Red Queen dynamics may be more generic than previously thought and, like our system, likely stable [45].

Previous theory work on the evolution of diversity via competition for discrete, resources has also supported the notion that evolution often drives ecosystems to remain in an unsaturated state via a common limiting resource [46] or a “diversification-selection balance” with substitutable resources [47]. While these unsaturated states are maintained by different processes than the low diversity metastable states presented here, the diversity of mechanisms promoting low diversity ecosystems hints toward their possible generality in nature.

Perhaps most intriguingly, low diversity states were actually further stabilized by small population sizes. In the individual-based model, reducing the environmental carrying capacity led to smaller population sizes as expected but also a smaller number of phenotypic clusters. For a given value of *K_max_* there exists some approximate maximal diversity that can stably exist. This remained true even when mutation size was increased. Increasing diversity with increasing ecosystem productivity has also been confirmed empirically [48]. In the adaptive dynamics simulations, increased mutation size allowed for mutants to jump into areas of positive invasion fitness, causing diversification. In the individual-based simulations this remained true, but these increased levels of diversity would not remain long, with one or more of the species dying out due to small population size stochasticity and increased competition. This means that in small populations, Red Queen dynamics were often perpetuated despite the presence of a higher diversity, globally stable ESS predicted by the adaptive dynamics. These non-equilibrium communities therefore remained in low diversity ESCs for perpetuity.

We should note that increased mutation size can also be considered as a proxy for migration between communities. The lack of a theoretical work on the interplay between eco-evolutionary and metacommunity dynamics is a known problem [49]. Experimental work measuring diversity as a function of immigration history finds that priority effects can play a significant role is shaping resulting communities [48, 50–53]. As the resulting communities are shaped by historical contingencies, this supports the idea that ecosystems may have many locally stable ESSs or limit cycles. Classical metacommunity theory suggests that community similarity will increase with high rates of migration between communities and low total population sizes [54]. Both of these factors proved analogous to our results. Increased mutation size modeling migration allowed communities to escape locally stable ESCs and converge on the unique globally stable ESS or limit cycle. Decreased carrying capacity leading to reduced maximal diversity forces all communities into fewer choices of metastable diversity patterns and therefore increased community similarity.

Niehus et al. [55] propose a model in which horizontal gene transfer acts in accordance with migration to homogenize microbial communities. While we find this idea compelling, we feel the complexity of HGT and immense diversity in microbial systems necessitate further investigation on the micro-evolutionary dynamics of HGT, representing an important and opportune area for future research.

Ultimately, the model and results presented here are broadly applicable to understanding how frequency-dependent selection can shape the realized diversity of natural communities and the dynamics of adaptive radiations. The pervasive presence of locally stable states or limit cycles in the adaptive dynamics and their further stabilization by finite population sizes suggests at their ubiquity in nature.

## Supporting information

### S1 Appendix. Full Model Details

We consider a general model of logistic growth and frequency-dependent competition based on a *d*-dimensional phenotype [17,18]. For simplicity the model does not consider spatial interactions. Evolution is modeled in three ways: adaptive dynamics, an individual-based model, and partial differential equations.

Ecological dynamics follow logistic growth governed by a carrying capacity 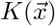 and a competition function 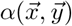 where 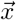 and 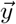 are the phenotypes of competing types with 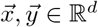.

The competition function is defined such that 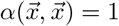, and for symmetric competition 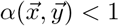 for 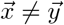. Thus, the Gaussian part of competition between types diminishes as their phenotypes becomes more distant and is maximal between individuals with the same phenotype. For symmetric competition the competition function is strictly Gaussian.

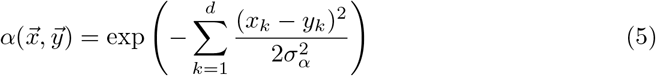

For asymmetric competition, the competition function also includes an additional term.

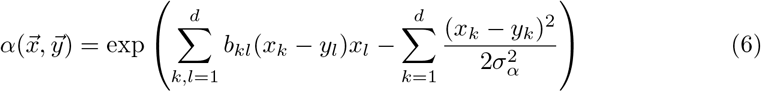

This first term is the first order of Taylor expansion of a higher order, non-symmetric competition function and thus represents the simplest form of adding non-symmetric dynamics to the Gaussian function [17]. It includes the coefficients *b_kl_*. For the non-symmetric competition simulations discussed here, a specific set of b coefficients were chosen that resulted in periodic evolutionary dynamics [provided in S5D Table]. The models were also run with many other values for these coefficients, including a survey of randomly chosen values, but results were qualitatively similar to the values chosen. For a review of how these values govern evolutionary dynamics in higher dimensions, please see Doebeli et al. [18].

The carrying capacity function 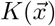 represents the equilibrium population size of a population of only individuals with phenotype 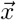. Two carrying capacity functions are discussed here. What is the quartic carrying capacity is defined as such.

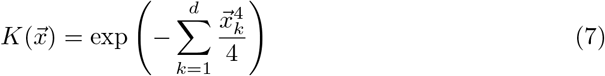

The radially symmetric carrying capacity is as follows.

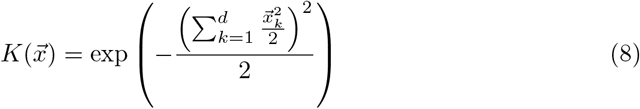

Both functions are similar in that they are maximal at the origin and of the fourth order. This ensures that there is stabilizing selection toward 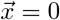 and that the phenotype space that is viable is bounded near the origin. The quartic carrying capacity function has an approximately square peak, while the radially symmetric function is circular [Fig. 1].

Together, ecological dynamics are as follows,

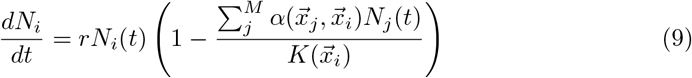

with *N_i_* representing the population size of individuals with phenotype 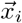, in a total population of *M* different phenotypes, and an intrinsic growth rate of *r*.

Using adaptive dynamics [21,25,26], the evolution of a phenotype 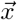 can be described by a system of differential equations 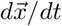. This is derived from the invasion fitness 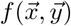, which is the per capita growth rate of a rare mutant with phenotype 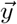 in a resident population with phenotype 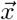.

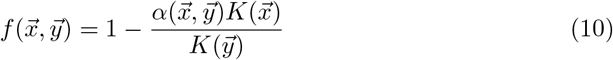

This invasion fitness function relates how the growth rate of the resident is always decreased with the introduction of a new mutant unless 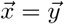 when 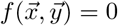. From this we can derive the selection gradient, *s*, as the derivative of the invasion fitness with respect to the mutant, 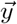, when it is equal to the resident, 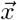.

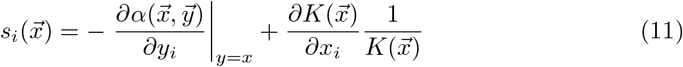

The adaptive dynamics are defined as

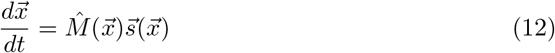

where 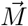 is the mutation-covariance matrix describing the rate, size, and covariance of mutations in each phenotypic dimension. For simplicity we assume this is the identity matrix. Any formulation of this matrix with a positive diagonal would only change the speed of the evolution in the different dimensions, but would not change the characteristics of any evolutionary dynamics or stable states. The adaptive dynamics are therefore a set of differential equations that describe the evolutionary dynamics in *d* dimensions for a given set of phenotypes.

Of note, while the Gaussian part of the competition kernel *α* affects whether diversification can occur and multi-species dynamics, because the selection gradient is evaluated at 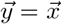, in the adaptive dynamics of monomorphic populations the Gaussian part of *α* disappears and we are left with just the effects of asymmetric competition. Because of the exponential nature of carrying capacity functions, the second term of the selections gradient, 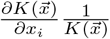, reduces to just the partial derivative of the inner function with respect to the resident. For the quartic case the adaptive dynamics thus as follows

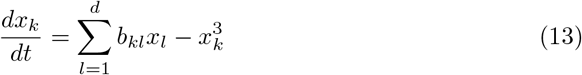

For the radially symmetric case the adaptive dynamics are

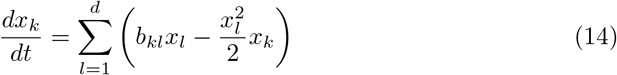

In order to include extinction and speciation events, an iterative algorithm is used: (1) solve for the ecological dynamics; (2) remove any newly extinct populations that fall below a minimum viable population size; (3) solve the adaptive dynamics for a given length of evolutionary time; (4) introduce a new mutant, with a phenotype a small, fixed distance from one of the resident populations or with a phenotype chosen from a Gaussian distribution with mean equal to the phenotype as one of the resident populations and a given variance 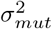; (5) remove the mutant if it is not ecologically viable (invasion fitness of the mutant is negative); and (6) repeat the process until either an evolutionary stable state or a given amount of time is reached.

At given intervals, clusters are calculated using a hierarchical clustering algorithm such that any two population with phenotypes within a small distance, *z_small_*, from each other are part of one cluster. Clusters are just an accounting device and do not affect dynamics. All parameters used can be found in S5A Table.

The individual-based model uses the same ecological dynamics as described above and simulated based on the Gillespie algorithm [57]. Individual birth rates are assumed to be constant and equal to 1, while death rates, *δ*, are frequency dependent and derived from the ecological dynamics.

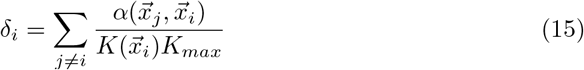

Here the competition function, *α*, and the carrying capacity function, *K*, are the same as in the adaptive dynamics, while *K_max_* is a constant that controls the height of the carrying capacity function to convert it from continuous to discrete populations. These rates are taken directly from the Lotka-Volterra equations, where the per capita growth rate of 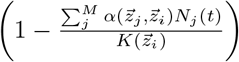 can be considered the birth rate minus the death rate. The individual-based model is thus a direct analog of the adaptive dynamics. For a larger discussion on the derivation of the individual-based dynamics of this model please refer to [12,31,33].

The simulation algorithm is as follows: (1) initiate the population with a randomly chosen initial population of a predetermined size; (2) update or calculate all individual death rates; (3) calculate the sum of all birth and death rates, 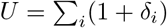; (4) increment time by a random amount drawn from an exponential distribution with mean equal to 1/*U*; (5) chose a random birth or death with probability equal to the ratio of the rate and the sum of all rates; (6) if a death rate is chosen, remove the chosen individual; if a birth rate is chosen, create a new individual with phenotype chosen from a Gaussian distribution with mean equal to the parent phenotype and variance 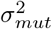; (7) rerun steps 2-6 until the population is extinct or a specified time is reached.

Phenotypic clusters are calculated using a hierarchical clustering algorithm in which every individual in a cluster has a phenotype within a small distance of at least one other individual in the cluster. Individuals are added to a cluster at birth or a new cluster is created if the individual does not fit in any existing cluster. A cluster is updated when a member individual dies to see if the cluster splits. All parameters used can be found in S5B Table.

### S2 Appendix. Numerical stability analysis

While we were unable to analytically determine the stability of the communities that arose in our simulations, we were able to test for evolutionary metastability numerically. However, this method is computationally expensive, so we were only able to test for stability on a small number of representative simulations.

To test the stability the final population in a simulation we extensively sampled random mutants around every resident in the community (250 mutants per resident). For each new mutant, we continued the adaptive dynamics simulation for a significant period of time (5% the length of the original simulation), while not allowing any other branching mutations to arise. We then checked whether the original residents and the mutant were all able to survive (*N_i_* > *N_small_*) and the mutant was able to diversify 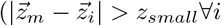 for a mutant with phenotype *z_m_* and residents *z_i_*) from the resident population. Any simulation in which no mutant can invade, coexist, and diversify is deemed an evolutionarily stable community.

While this method is unable to definitively prove stability as it is always possible that an extremely rare mutant could diversify or drive multiple residents extinct, we can at least demonstrate the metastability (as defined in the main text; any state that is maintained for a period significantly longer than its convergence) of these communities. An illustration of metastability can be seen in Figure 11. Here, the convergence to the metastable cycle with 10 species takes approximately 25 evolutionary time units, after which the system resides in the metastable state until the end of the simulation, which lasts for another 975 time units. A clearer depiction of the early convergent dynamics can be seen in the main text in Figure 3.

**Fig 11.**
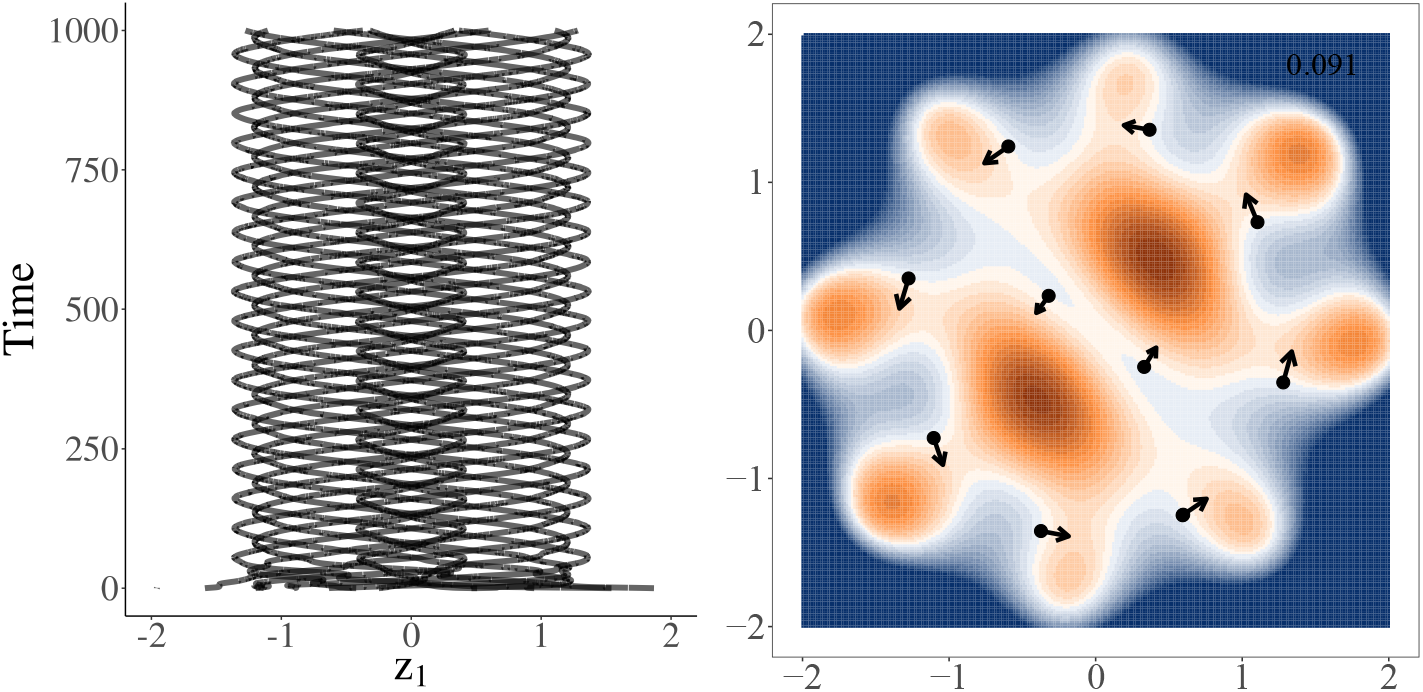
Convergence to metastable limit cycle. Here we define metastability operationally as any system that resides in a state much longer than it took to converge to it. Here, the system converges to a stable limit cycle in approximately 25 time steps and remains there for the duration of the simulation. The simulation were run using the radially symmetric carrying capacity and initiated with 18 random species. Panel A shows the complete history of evolutionary dynamics. Panel B is a depiction of the population at the end of the simulation. Colors in panel B represent the invasion fitness (per capita growth rate of a new mutant if it were to arise). Positive invasion fitness is shown in shades of orange (maximum of 0.091), negative in blue, and invasion fitness equal to zero in white. Arrows are proportional to the square root of the selection gradient for each species.

This numerical stability analysis was used to test each simulation illustrated in the main text (Figs 2 and 4). Using the same small mutations as the original simulations, not a single mutant (of over 30,000 tested) was able to invade, coexist with, and differentiate from the original community. The same was true when we instead tested Gaussian mutations (*σ_mut_* = 0.05). Only when we tested larger mutations (Gaussian mutations with *σ_mut_* = 0.1) were 7 mutants (0.02% of those attempted) from two different simulations able to break the stability and diversify the community into a new, higher diversity state.

At least in the representative simulations tested, only large or extremely rare mutants are able to shift the community, proving that these states are indeed evolutionarily metastable.

Figures from the numerical stability analysis for each of these simulations can be found on-line at https://www.zoology.ubc.ca/~rubin/AltEvoDiversity/.

### S3 Appendix. PDE model

In addition to the adaptive dynamics and individual-based models, we also ran partial differential equation simulations. PDEs represent the infinite population approximation of individual-based models [31, 33] and are useful for relaxing the phenotypes represented as a delta function and the separation of timescale between ecology and evolution assumptions of adaptive dynamics without the stochasticity of individual-based models.

The large population limit of the individual based model gives the deterministic formulation of partial differential equations.

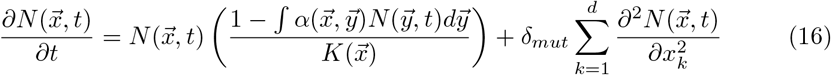

Here, 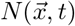 is distribution of the population with phenotype 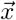 at time *t* and *δ_mut_* is diffusion coefficient that can be thought of as analogous to the rate of mutations and will not have qualitative effect on our simulations. The competition and carrying capacity functions are the same as described above. The PDE is numerically solved over a lattice. Because of memory limitations, using a high resolution lattice is infeasible, though we were able to run all 2-dimensional PDE simulations on a 200 x 200 lattice. This lattice is more than detailed enough to reveal any peaks and patterns that appear. The local maxima of the resulting distributions represent centers of clusters of individuals in the individual-based model.

Notably, like the individual-based model, the PDEs are not restricted to approximation of adaptive dynamics that phenotypes are represented as delta functions. Instead, the distributions produced by the PDEs “take up space” in the phenotype space. This can result in configurations with fewer local maxima than adaptive dynamics populations when run with the same parameters. Additionally, because the PDEs are a diffusion process, configurations cannot be trapped by small mutations. This makes the PDE simulations unsuited to studying the presence or absence of alternative stable configurations. They are however an useful comparison to the adaptive dynamics and individual-based models that are fully deterministic and free of the assumptions intrinsic to adaptive dynamics.

When PDE simulations are run with symmetric competition, the resulting population densities are configured in the same 4×4 grid and two concentric circles as the comparable adaptive dynamics simulations (Figs 2, 12). The same concentric circles appear as ridges of high population density, but only the inner is peaked and far less distinctly than with the quartic grid. As the adaptive dynamics assumes species can be represented by a single phenotype without variance, continuous diversification around the circles is the only way for the population to mimic the circular ridge predicted by the PDE. This implies that with infinite population size (adaptive dynamics and PDE), simulations with symmetric competition and a radially symmetric carrying capacity eventually (though this process is slow) result in no distinct species or phenotypic clusters. However, as noted in the main text, this situation of continual diversification along concentric circles in trait space is likely a degenerate case that is cause by a radially symmetric carrying capacity function and Gaussian competition.

**Fig 12.**
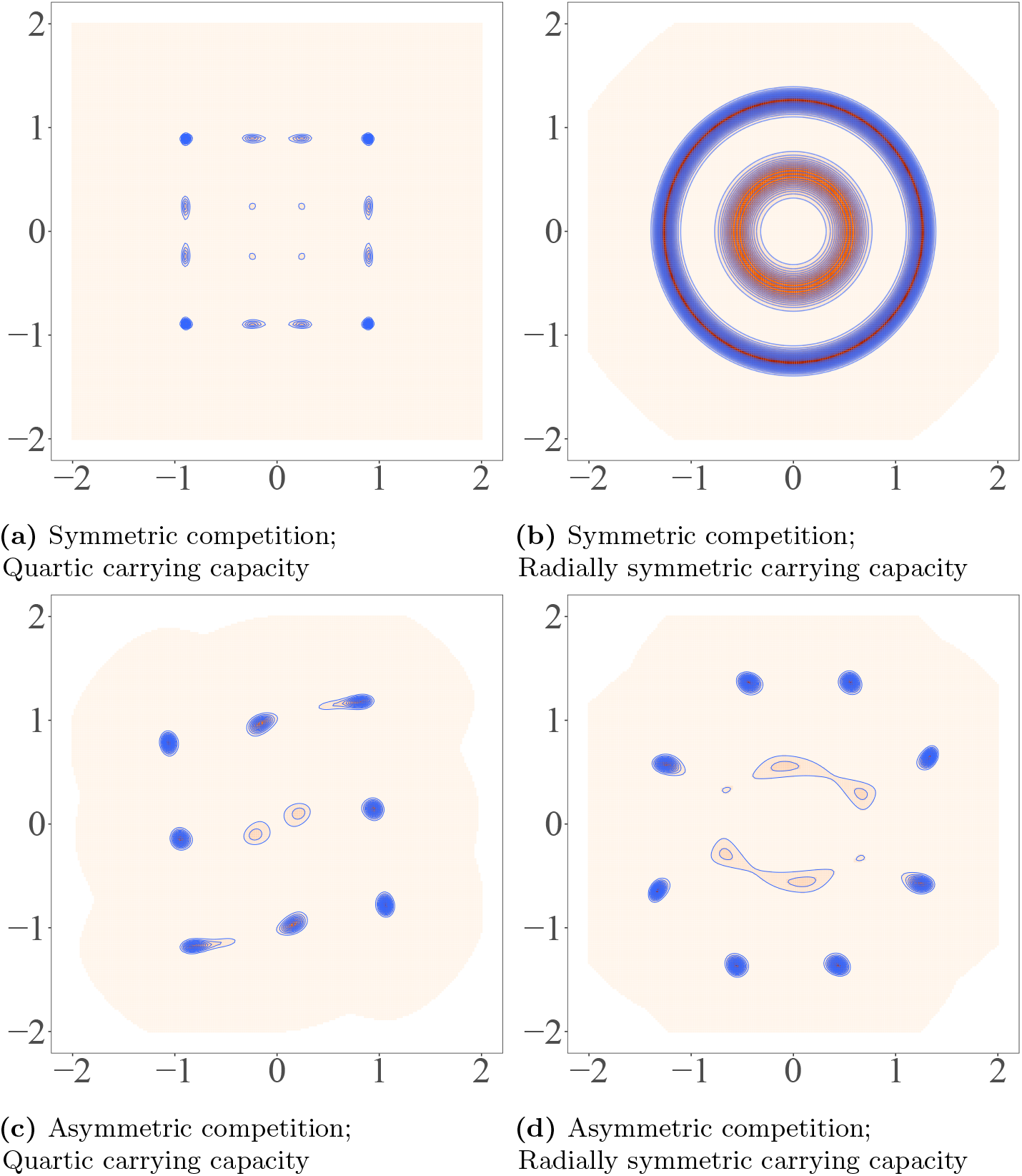
The final population density when simulated using a PDE. Higher densities are darker and lower densities are lighter. Contour lines are shown in blue. 3-dimensional, interactive versions of the same figures and videos of the dynamics are available on-line (S4 Video). Population distributions generated using the PDE models result in similar configurations to those emergent from the Adaptive Dynamics, but with fewer distinct species resulting from intra-species phenotypic variance. The asymmetric competition and radially symmetric carrying capacity simulation exhibits the same clockwise rotation displayed in the adaptive dynamics and individual-based simulations.

For the asymmetric competition, PDE simulations with both quartic and radially symmetric carrying capacities result in similar, but less diverse patterns in comparison to the highest evolutionary stable diversity state from the adaptive dynamics simulations (10 versus 14 phenotypes for the quartic carrying capacity and 14 versus 16 phenotypes for the radially symmetric case) (Figs 4, 12). The reduced diversity is likely because the peaks in the adaptive dynamics are delta peaks, while those in the PDE have non-zero variance in phenotype space, restricting the number of distinct peaks. The radially symmetric carrying capacity simulation did exhibit the same clockwise rotation in both rings as displayed in the adaptive dynamics and individual-based models (video available in on-line, S4 Video).

There has been previous theoretical work that predicts variation within species clusters has a negative effect on coexistence between competing species [56]. As species now occupy a distribution in trait space, rather than a single point, niche differentiation between species is reduced, impeding the maintenance of higher numbers of distinct phenotypes. As the PDE is the infinite population limit of the individual-based models, we would expect the same lower diversity at the global ESC when the individual-based simulations are run with very large communities and mutation sizes greater than some *ε* (these simulations were computationally infeasible to run). While we don’t expect the other results (the presence of locally stable ESSs or limit cycles) from this paper to be affected by the lower diversity, we feel it is a salient point worth considering when comparing theoretical models of diversification to expectations for natural populations.

All parameters used can be found in S5C Table.

### S4 Video. Videos and additional figures

Videos of the evolutionary dynamics for all simulations highlighted in the paper (including adaptive dynamics, individual-based, partial differential equation, and stability analysis simulations) can be found on-line at https://www.zoology.ubc.ca/~rubin/AltEvoDiversity/. Additional figures including 3-dimensional, interactive landscapes of the final population density of PDE simulations can be found here as well.

### S5A Table. Default parameters for adaptive dynamics simulations

**Table 1.** Parameters and variables used to generate adaptive dynamics data and figures.

| Parameter | Value | Definition |
| --- | --- | --- |
| $\sigma_\alpha$ | 0.5 | Standard deviation of the competition kernel |
| $r$ | 1.1 | The intrinsic growth rate |
| $\epsilon_{mut}$ | 0.005 | Mutation size |
| $T_{max}$ | $10^5$ | Number of time steps simulations run for |
| $T_{split}$ | 0.2 | Time between each branching attempt |
| $v_{min}$ | $10^{-10}$ | Evolutionary velocity* deemed stable to stop simulation early |
| $N_{small}$ | $10^{-8}$ | Population density which populations are considered extinct |
| $z_{small}$ | $10^{-3}$ | Phenotypic distance two species are deemed identical and merged |

* Evolutionary velocity is defined as the population weighted average magnitude of the rates of the phenotype vectors:

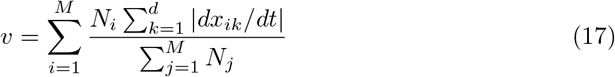

After each step solving the adaptive dynamics (step 3 as outlined above), the evolutionary velocity is checked, and if *υ* < *υ_min_*, the simulation is deemed stable and stopped.

### S5B Table. Default parameters for individual-based simulations

### S5C Table. Default parameters for partial differential equation based simulations

### S5D Table. Asymmetric competition parameters

The *b* values dictating the competition asymmetry used in all asymmetric competition simulations are below.

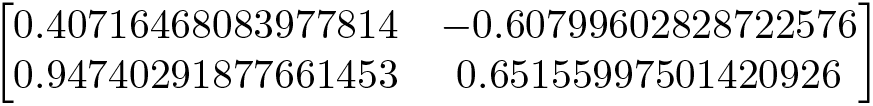

**Table 2.** Parameters and variables used to generate individual-based model data and figures.

| Parameter | Value | Definition |
| --- | --- | --- |
| $\sigma_\alpha$ | 0.5 | Standard deviation of the competition kernel |
| $r$ | 1.1 | The intrinsic growth rate |
| $\sigma_{mut}$ | 0.005 | Standard deviation of mutation |
| $T_{max}$ | $10^5$ | Number of time steps simulations run for |
| $K_{max}$ | 400 | The height of the carrying capacity function at the origin |
| $\delta_{cluster}$ | $50\sigma_{mut}$ | The maximum distance between individuals of the same cluster |

**Table 3.** Parameters and variables used to generate partial differential equation model data and figures.

| Parameter | Value | Definition |
| --- | --- | --- |
| $\sigma_\alpha$ | 0.5 | Standard deviation of the competition kernel |
| $N_{bins}$ | 200 | The number of bins in each dimension |
| $z_{range}$ | $(-2, 2)$ | The minimum and maximum phenotype in each dimension |
| $\delta_{mut}$ | 0.01 | the amplitude of diffusion modeling mutation |
| $T_{max}$ | $10^3$ | Number of time steps simulations run for |
| $\sigma_N$ | 0.1 | Standard deviation of the initial spread around the origin |
| $\delta_t$ | 2 | The integration time step |

## Acknowledgments

II acknowledges support from FONDECYT project 1200708. MD acknowledges support from NSERC, grant no. 219930.

## Notes

### Competing Interest Statement

The authors have declared no competing interest.

